# Experimental estimates of germline mutation rate in eukaryotes: a phylogenetic meta-analysis

**DOI:** 10.1101/2023.01.24.525323

**Authors:** Yiguan Wang, Darren J. Obbard

**Affiliations:** Institute of Ecology and Evolution, University of Edinburgh, Charlotte Auerbach Road, Edinburgh EH9 3FL, UK

**Keywords:** de novo mutation rate, eukaryotes, indels, phylogeny

## Abstract

Mutation is the ultimate source of all genetic variation, and over the last ten years the ready availability of whole-genome sequencing has permitted direct estimation of mutation rate for many non-model species across the tree of life. In this meta-analysis we make a comprehensive search of the literature for mutation rate estimates in eukaryotes, identifying 140 mutation accumulation (MA) and parent-offspring (PO) sequencing studies covering 134 species. Based on these data, we revisit differences in single nucleotide mutation (SNM) rate between different phylogenetic lineages and update the known relationships between mutation rate and generation time, genome size, and nucleotide diversity—while accounting for phylogenetic non-independence. We do not find a significant difference between MA and PO in estimated mutation rates, but we confirm that mammal and plant lineages have higher mutation rates than arthropods, and that unicellular eukaryotes have the lowest mutation rates. We find that mutation rates are higher in species with longer generation times and larger genome sizes, even when accounting for phylogenetic relationships. Moreover, although nucleotide diversity is positively correlated with mutation rate, the gradient of the relationship is significantly less than one (on a logarithmic scale), consistent with higher mutation rates in populations with smaller effective size. For the 29 species for which data are available, we find that indel mutation rates are positively correlated with nucleotide mutation rates, and that short deletions are generally more common than short insertions. Nevertheless, despite recent progress, no estimates of either SNM or indel mutation rates are available for the majority of deeply-branching eukaryotic lineages—or even for most animal phyla. Even among charismatic megafauna, experimental mutation rate estimates remain unknown for amphibia and scarce for reptiles and fish.

**Lay Summary:** Over the past decade, the sequencing revolution has led to an ever-increasing number of mutation-rate estimates from mutation accumulation or parent-offspring sequencing studies in eukaryotes. However, studies rarely quantify to what extent the mutation rate varies among these species. Also, despite strong predictions as to how mutation rate should vary with (e.g.) generation time, there have been few recent or wide-ranging analyses of such predictors while accounting for the inherent similarity between closely-related species. Of particular note, there has been surprisingly little effort to robustly test the ‘drift barrier’ hypothesis that mutation rates should decrease with increasing effective population size. In this study, we used a comprehensive literature search to identify all the available experimental estimates of mutation rate in eukaryotes and subject them to phylogenetic mixed-model analyses. We find that per-nucleotide per-generation mutation rates differ by orders of magnitude among species: plants and mammals tend to have higher mutation rates than arthropods, and unicellular organisms have the lowest mutation rates. Our analysis also shows that mutation rates increase significantly with increasing generation time and genome size, and nucleotide diversity increases with mutation rate with a gradient less than one—as predicted by the drift-barrier hypothesis.

## Introduction

The per-nucleotide per-generation de novo mutation rate, µ, is a key parameter in population and evolutionary genetics, appearing either alone or combined with effective population size (N_e_) in the compound parameter θ. However, it seems implicit that µ is of no interest to many evolutionary studies; researchers often use µ from one species when analysing another—as if µ were exchangeable in a way that N_e_ is not (e.g. Wilding, 2017). This is partly because the historically high cost of sequencing made estimates prohibitively expensive, but may also be because variation in µ among species is widely thought to be negligible compared to variation in N_e_. But, if the mutation rate varied as widely among multicellular eukaryotes as many other traits, there would be much less need to invoke variation in N_e_ when explaining variation in (e.g.) genetic diversity. One reason why µ may vary relatively little is if it is minimised to a lower limit imposed by drift: the ‘drift barrier’ hypothesis (Lynch, 2008, 2010). However, it might be that in reality mutation rate is optimised to well above the drift-barrier limit by selection in favour of adaptability or the cost of replication fidelity (Kimura, 1967; Liu et al., 2021; Peck et al., 2018). If this were the case, then we might find that mutation rates varied dramatically, even among closely related taxa.

Early estimates of the mutation rate leveraged visible phenotypes, such as the occurrence of genetic diseases in humans (Nute et al., 1984; Trimble et al., 1974). For example, using haemoglobin M disease Stamatoyannopoulos et al. (1982) estimated the mutation rate in the human haemoglobin beta gene at 7.4 × 10 per bp per generation. However, such estimates are prone to bias, for example if the mutation is required to be autosomal dominant and non-lethal (Nute et al., 1984), and such approaches are not easily applicable in organisms for which disease is hard to observe. More commonly, using the equivalence between mutation and neutral substitution rates (Kimura, 1968), many studies have used time-calibrated sequence divergence to estimate µ. For example, comparison between humans and chimpanzees gives rise to estimates on the order of 10 per bp per generation (Drake et al., 1998; Kondrashov et al., 1993; Nachman et al., 2000). However, such indirect estimates are also limited (Kondrashov, 2003). First, because generation time and calibration dates are prone to substantial uncertainty (e.g. Obbard et al., 2012). Second, because it is hard to distinguish between unconstrained and weakly constrained sites (Harmon et al., 2021). Third, because mutation saturation in hotspots may lead to an underestimation, especially for sequences with a long divergence time (Sigurðardóttir et al., 2000). In addition, phylogenetic-calibration approaches reflect the long-term mutation rate, averaging across biologically important factors such as generation time, sex differences, life-stage differences, and inter-individual variation.

In contrast to phenotypic and phylogenetic approaches, mutation-accumulation (MA) and parent-offspring (PO) sequencing can provide direct estimates of the per-generation mutation rate. Mutation-accumulation utilises the accumulation of spontaneous mutations in a single inbred or asexual genome over multiple generations, by minimising the effectiveness of natural selection (Halligan et al., 2009). Given the low mutation rates in eukaryotes, the multiple generations in an MA experiment allow sufficient mutations for them to be detected at the end. Although the first use of MA to estimate µ phenotypically dates back to a study by Mukai et al. (1977), the availability of whole genome sequencing and its decreasing price have made sequencing approaches more widely applicable.

Mutation accumulation best suits easily-maintained species with short generation times, for example nematodes (Denver et al., 2004; Konrad et al., 2019; Meier et al., 2014; Saxena et al., 2019), fruit flies (Haag-Liautard et al., 2007; Huang et al., 2016; Keightley et al., 2009; Schrider et al., 2013), green algae (Lopez-Cortegano et al., 2021; Ness et al., 2012), and water fleas (Ho et al., 2020; Keith et al., 2021). However, MA may underestimate mutation rates (Lynch et al., 1998) because natural selection will still come into play for mutations that are very highly deleterious, such as those that cause complete sterility or are lethal (Baer et al., 2007).

By sequencing parents and offspring, and counting the mutations arising over a single generation, the parent-offspring approach avoids purging deleterious mutations, except those that are dominant lethal, and so provides the most direct estimate of mutation rate. It is not limited to model species (Keightley et al., 2014; Yang et al., 2015), and is suitable for large animals outside of the laboratory (Harland et al., 2017; Wang et al., 2022). Parent-offspring sequencing has been particularly widely used in humans, where MA is not applicable. However, the challenge of robustly identifying new mutations in natural heterozygous genomes is formidable, as the number of mutations arising within one generation is small when compared to the impact of sequencing/mapping errors and the genetic variation of heterozygotes. For example, Bergeron et al. (2022) compared the estimates from five different research groups for the same family of rhesus macaques and found that the highest estimates could be twice the lowest ones. Minimising the rates of false positive and false negative mutation-calls is therefore critical for the PO approach (Yoder et al., 2021).

Relaxed-clock phylogenetic studies generally indicate there is substantial among-lineage variation in the evolutionary rate, suggestive of variation in mutation rates among taxa (Ho et al., 2015). But such studies cannot separate the impact of generation time and mutation rate, and may be biased by variation in the action of (weak) selection (Harmon et al., 2021). As more direct experimental estimates have become available, it has become possible to draw comparisons within particular clades such as primates (Chintalapati et al., 2020), or vertebrates (Bergeron et al., 2022; Bergeron et al., 2023; Yoder et al., 2021). However, there has not been a wider analysis, nor one that includes both PO and MA studies.

In addition, many previous studies have not taken an explicitly phylogenetic analytical approach (e.g. Bergeron et al., 2022; Katju et al., 2019; Yoder et al., 2021). This could in principle lead to pseudoreplication in the analyses that is caused by the non-independence among taxa that arises from their shared ancestry (reviewed in Freckleton, 2009). However, this problem can be mitigated by explicitly modelling covariance among species that is contributed by their patterns of relationship, and generalised linear mixed modelling approaches are available to do this (Hadfield et al., 2010; Halliwell et al., 2022; Lynch, 1991).

Here we perform a comprehensive phylogenetic meta-analysis of published eukaryotic single nucleotide mutation (SNM) and insertion-deletion (indel) mutation-rate estimates that use direct sequencing of multiple generations, either through MA or PO comparison. We focus on the variation in mutation rate among species, and the relationship between mutation rate estimates and the experimental method, generation time, genome size, and genetic diversity.

### Results and Discussion Studies included

We searched for eukaryotic mutation-rate studies in Clarivate Web of Science published prior to 21 September 2022, and then we manually examined these studies and identified those containing mutation rate estimates (see Methods). For the majority of our analyses we also included data from one further recent study of 68 vertebrate species by Bergeron et al. (2023), published after our initial submission. In total, we included 140 studies using either MA or PO approaches, covering 134 species (see Availability of data and materials). Studies on primate mutation rate accounted for over a third of this literature, and 82% (45/55) of the primate studies were on humans. Arthropods accounted for 17% of the identified literature, followed by fungi (11%), plants (8%), non-primate mammals (7%), and nematodes (5%) (Fig. 1A). We only identified four studies on fish, two studies on birds and one study on reptiles. Across all groups, most studies focused on model species: yeast in fungi, Drosophila in arthropods, mice in mammals, and Arabidopsis in plants. These estimates tended to derive from MA analyses rather than PO studies (61 vs. 43 among non-human studies). However, as sequencing has become cheaper, there has been a rapid increase in the number of published experimental estimates of the mutation rate, with an average of 11 studies each year in eukaryotic species between 2010 and 2022 (Fig. 1B).

**Figure 1:**
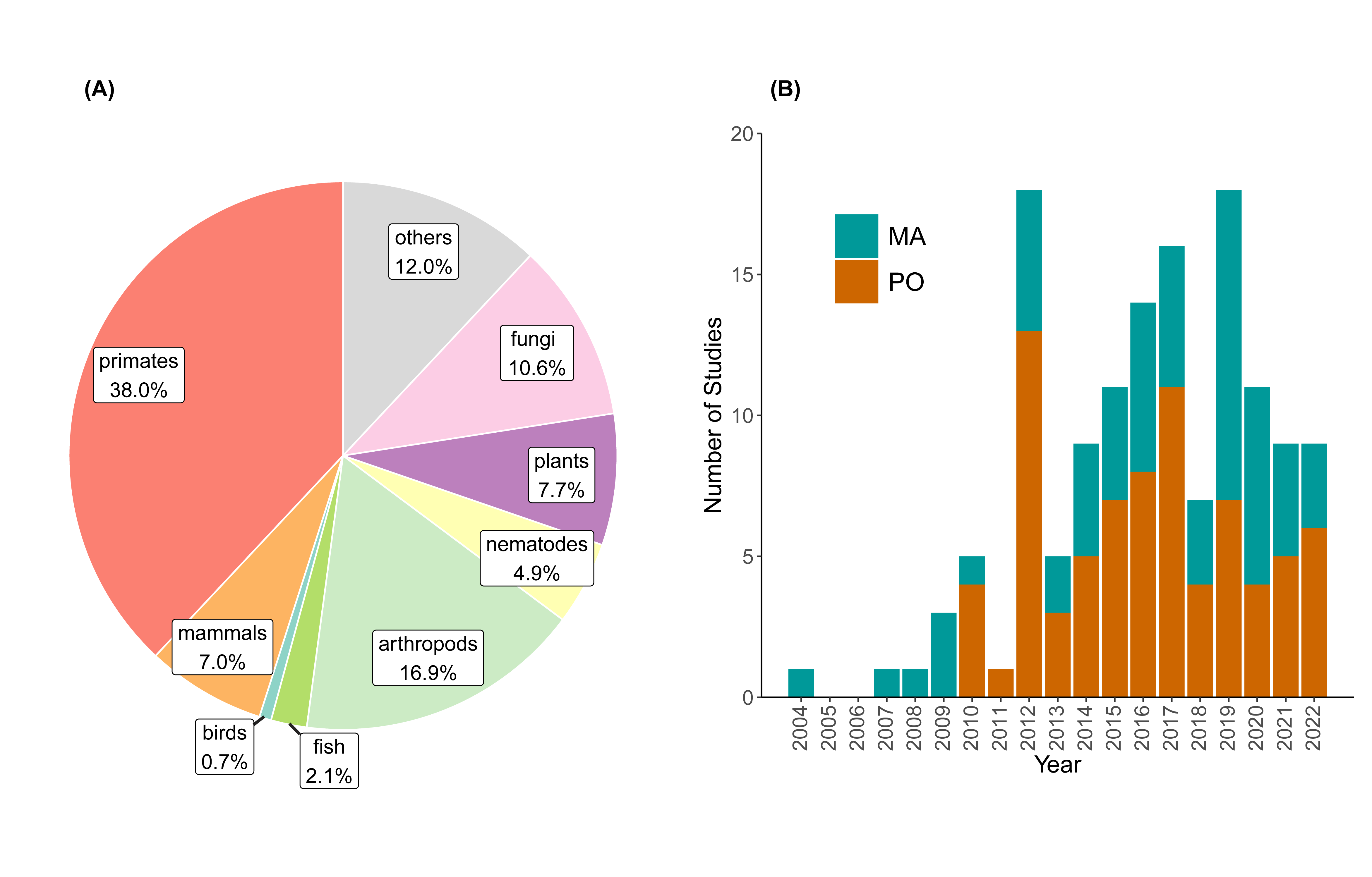
A summary of de novo mutation rate studies published up to 21 September 2022. (A) The proportion of studies in different taxonomic groups. (B) The number of studies by year. MA: mutation accumulation studies, PO: parent-offspring studies.

### Single nucleotide mutation rates across eukaryotes

To summarise the single nucleotide mutation rates in different lineages, we obtained a time-scaled phylogenetic tree of all represented species from http://www.timetree.org/ (Kumar et al., 2022), and used this to fit the covariance among species in a Phylogenetic Generalised Linear Mixed Model (PGLMM) using the Bayesian mixed model package MCMCglmm (Hadfield et al., 2010). Treating species as a random effect, this provides a posterior estimate of mutation rate for each species, given the observations and relationships to other species (see Methods). In these data, the phylogenetic effect accounts for 96.4% of the random-effects variance, with a 95% HPD interval of 94.9% to 97.9%.

Among all the studied species, a ciliate (Tetrahymena thermophila) had the lowest mutation rate, estimated at 0.01 × 10 per generation per base pair, while a fungus (Marasmius oreades) had the highest rate, estimated at 55.58 × 10 —a difference of over five thousand-fold. The estimates for unicellular species were generally more than an order of magnitude lower than rates in multicellular species (below; Fig. 2), even compared to species in the same clade, e.g., fungi. This may either be attributable to there being a single cell division per generation, or possibly to a limit set by the drift barrier in species with larger N_e_ (Sung et al., 2012). We also estimated relatively low SNM rates for nematodes at 2.44 × 10, arthropods at 2.82 × 10^-9^, and fish at 5.55 × 10 . For plants, the clade root was predicted as 3.66 × 10, but a large difference was observed between duckweed and other plant species (0.22 × 10^-9^ vs. 19.8 × 10). The high mutation rates in most plant species might be attributable to the lack of a segregated germline (Burian, 2021; Cruzan et al., 2022; Lanfear, 2018; Wang et al., 2019). Mammals had moderate mutation rates with a root estimate at -9 8.36 × 10 and species (excluding primates) ranging from 5.64 × 10^-9^ in pigs to 11.68 × 10 in killer whales. Primates had higher mutation rates, with a root estimate at 9.37 × 10^-9^. Of note, as the most studied species, human mutation rate was estimated as 13.41 × 10 with 95% CI 11.61 to 15.28 × 10^-9^, which is also the highest among mammals. Similar to mammals, birds and reptiles had high mutation rate, with root estimates at 10.43 × 10^-9^ and 12.41 × 10^-9^, respectively.

**Figure 2.**
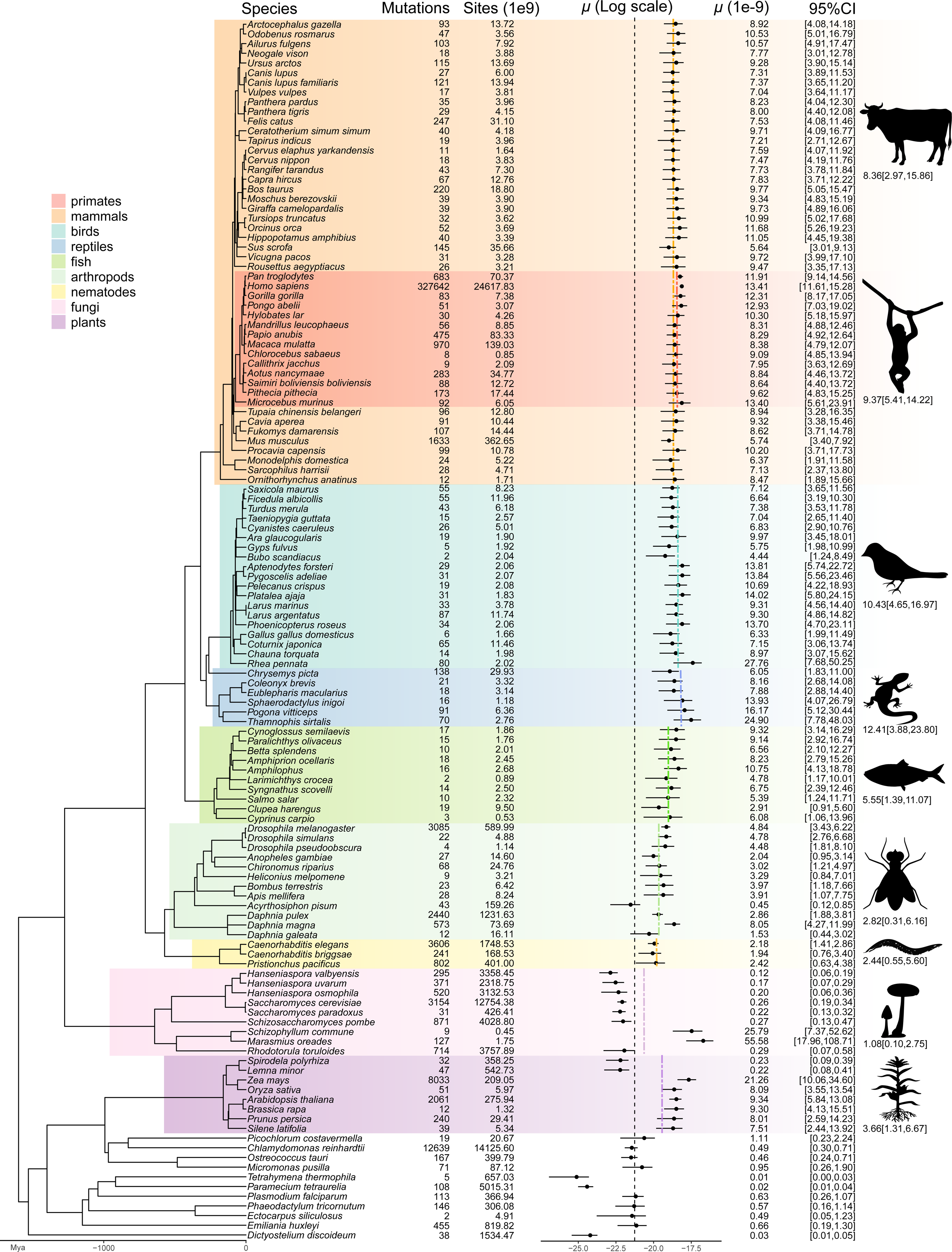
A phylogenetic meta-analysis of SNM rates for different species identified from the literature. Mutation rates and 95% CIs were estimated using a PGLMM. For each annotated clade, we estimated the rate for their most recent common ancestor (clades marked in colour; estimates in dashed lines and below taxon icons to the right). The vertical black dashed line indicates the mutation rate of common ancestor of the tree.

To assess a possible impact of the experimental method on rate estimates, we included the experimental approach (MA or PO) as a fixed effect in the PGLMM. Overall, we did not find a significant difference between the two methods (MCMC P = 0.155). We then examined comparisons between MA and PO in fruit flies and mice more closely, as these two species have been investigated using both methods. The rate estimated using MA in fruit flies was 5.28 × 10^-9^, but 3.20 × 10^-9^ using PO (Fig. S1), and in mice the MA estimate was 5.40 × 10^-9^ while the PO estimate was 4.05 × 10 (Fig. S2), and these differences were similarly not significant. Note, however, that the mutation rate may be heterogeneous among different populations: for D. melanogaster, the estimated mutation rate from African populations was lower than that from European population and North American population (Chan et al., 2012; Wang et al., 2023). To make a more robust analysis would require samples of the same population but using different methods (MA or PO), which are not yet available.

### Biological predictors of the SNM rate

Many biological factors have been proposed to be associated with mutation rate, including generation time (Bailey et al., 1991; Keightley et al., 2000; Li et al., 1987; Wu et al., 1985) and genome size (Drake, 1991; Lynch, 2010). To test for a dependence of mutation rate on these factors, we ran three further models: the first two including these factors each separately as an additional fixed effect in a PGLMM, and the last one including both of them simultaneously.

We found a significant positive relationship with generation time (Bayesian PGLMM MCMC P < 0.0001; Fig. 3A), such that a longer generation time predicted a higher per-site mutation rate per generation. If mutations resulted only from replication errors, and most species had a similar number of cell divisions per generation, then the per-generation mutation rate would not scale with generation time (Thomas et al., 2014; Wu et al., 1985). However, although few estimates are available, in reality the number of germline cell divisions per generation varies within and between species. In humans, there are around 401 cell divisions per 30-year generation in males and 31 in females (Drost et al., 1995; Ohno, 2019), but in mice it is 62 per 9-month generation in males and 25 in females. In Drosophila the sexes are much more similar, with an estimated 35.5 cell divisions in a 18-day generation for males but 36.5 in an 25-day generation in females in D. melanogaster (Drost et al., 1995). Within a species, a long generation time is likely to permit more cell divisions and thus a higher mutation rate per generation, and many studies in humans and primates have shown that mutations are heavily male-biased and correlated with paternal ages (Kaplanis et al., 2022; Kong et al., 2012; Thomas et al., 2018; Wang et al., 2020).

**Figure 3.**
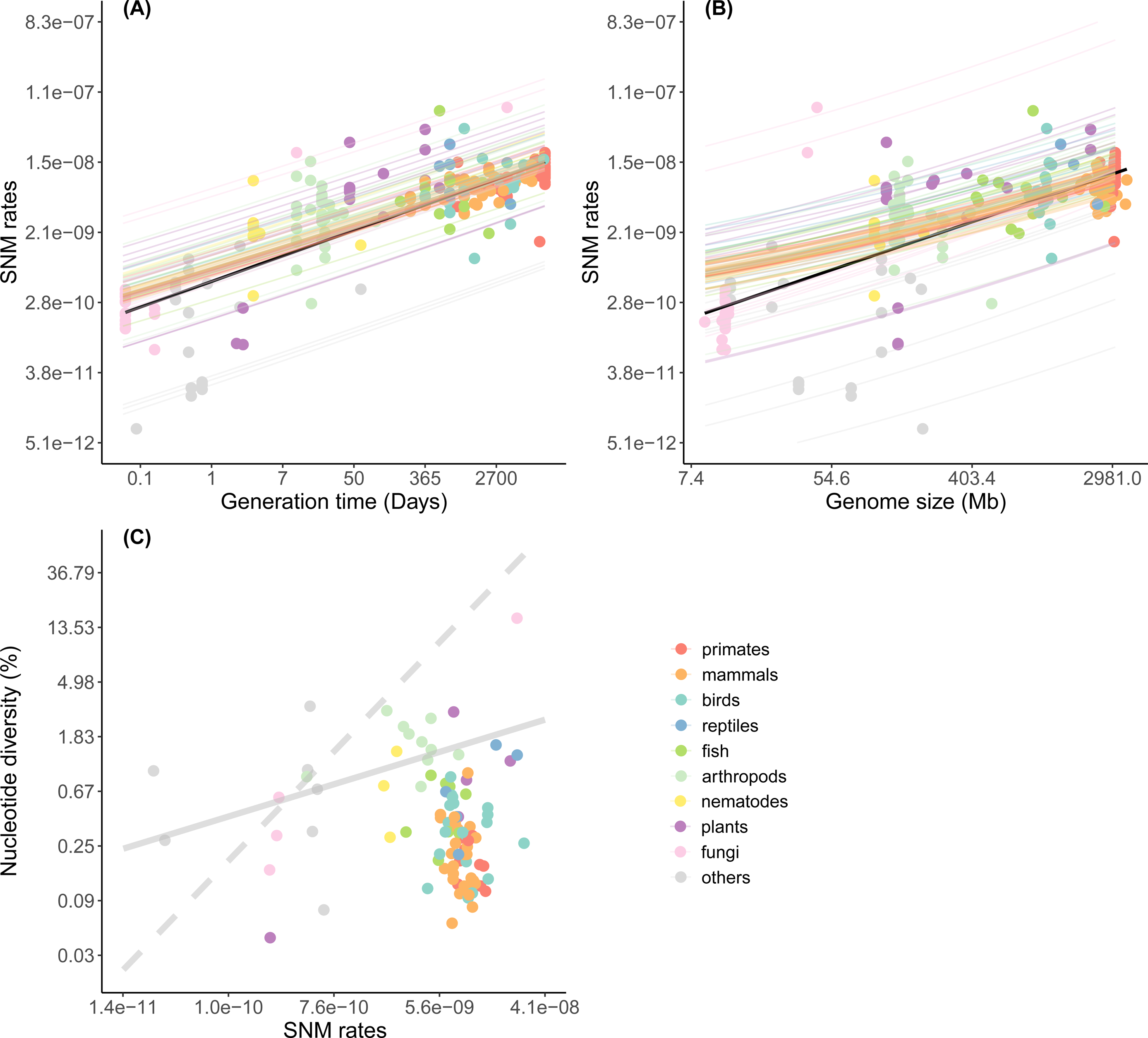
Phylogenetic linear regressions of SNM rate on (A) generation time for 116 species, (B) on genome size for 108 species. The solid black lines represent simple linear regressions; the coloured lines represent different PGLMM regressions conditional on each of the species random effects. (C) a regression of pairwise nucleotide diversity (π) on SNM rate (µ), fitted as a bivariate model in MCMCglmm for 102 species. The solid grey line indicates the model regression line; the dashed grey line indicates the expected slope for θ_π_ = 4N_e_μ on the log scale. All axes are plotted on a log scale.

Alternatively, mutations may not result predominantly from replication errors. Recent studies suggest that not all germline mutations track cell divisions, and DNA damage may contribute to the mutation rate (Wu et al., 2022; Wu et al., 2020). Independent of cell divisions, this mutation rate would be correlated with calendar time, and thus species with a longer generation time would have a higher rate. The relationship we observe between generation time and per-generation mutation rate could therefore be a consequence of either a greater number of cell divisions or of accumulating damage over time, and it will be hard to separate these potential causes given the limited data available on the number of germline cell divisions for different species (Gao et al., 2016; Wu et al., 2022).

We also found evidence that larger genome size predicts a higher per-generation mutation rate (MCMC P = 0.0020; Fig. 3B). Mutation rate has previously been found to correlate negatively with genome size in prokaryotes, a phenomenon termed “Drake’s rule” (Bourguignon et al., 2020; Drake, 1991; Marais et al., 2020), but positively in eukaryotes (Lynch, 2010).

In each of the two models above, we fitted generation time and genome size individually as predictors of mutation rate in a regression model. However, these factors might be highly correlated: large organisms tend to have longer generation times and often larger genomes (Buffalo, 2021; Martin et al., 1993), and is therefore hard to distinguish between casual relationships and confounding correlations. Including generation time and genome size together as fixed terms in a single model resulted in only generation time being significant with MCMC P < 0.001 (versus MCMC P = 0.301 for genome size).

### The drift-barrier hypothesis

The most direct test of the drift-barrier hypothesis would be a regression of SNM rate on N_e_. However, long-term N_e_ can only be estimated from genetic diversity (i.e. π) by making use of the relationship θ_π_ = 4N_e_μ, and any uncertainty in μ could induce a spurious correlation in a regression of μ on π/4μ. Instead, error in both π and μ can be accounted for by fitting a bivariate linear mixed model, parameterised in terms of a regression of π on μ (see Methods). Then, on a logarithmic scale, the gradient of this relationship would be expected to be 1 if μ and N_e_ are independent of each other, and lower than 1 if species with large N_e_ tended to have lower μ. Nevertheless, such a relationship could be driven by other factors. For example, if species with small N_e_ tend to have a longer generation time, and a longer generation time causes higher mutation rates (i.e. Fig. 3A), then a higher μ in species with low N_e_ could be driven by a mechanistic generation-time effect, rather than the drift barrier. We attempted to account for this by including generation time in our analysis as a predictor of both μ and π, thereby accounting for the direct impact of generation time on the relationship between them. In this bivariate phylogenetic model, μ increased with generation time (MCMC P < 0.001) whereas π did not (MCMC P = 0.501), and the gradient of the regression of π on μ was significantly less than 1 (gradient = 0.535, 95% CI [0.259, 0.832]; MCMC P = 0.004). This suggests that populations with larger N_e_ tend to have a lower mutation rate even after accounting for their shorter generation times—as predicted by the drift-barrier hypothesis (Lynch, 2010). Note that excluding generation time as a predictor resulted in an even stronger effect (gradient = 0.300 [0.098, 0.486]; MCMC P < 0.001; Fig. 3C), whereas removing unicellular species from the analysis (because of their low mutation rates) resulted in a gradient that was still much less than 1 (gradient = 0.439 [0.155, 0.746]; MCMC P < 0.001).

### The mutation rates of short indels across eukaryotes

Although the rate of indel mutation is much less commonly estimated than the SNM rate, our literature search identified estimates for 29 species. We found that the indel mutation rate across eukaryotes displayed a similar pattern to that of the SNM rate (Fig. 4), with a correlation coefficient between the two of 0.75 (95% HPD CI: 0.44, 0.98; MCMC P < 0.001) using PGLMM, with the SNM rate being generally higher than the indel mutation rate, except in two lineages (nematodes and amoebae) for which the indel mutation rate was apparently higher (Fig. 5A). However, the mutations in the nematode study (Denver et al., 2004) were specifically collected from mononucleotide repeats, which may artificially inflate the indel mutation rate estimate in that study (Konrad et al., 2019; Rajaei et al., 2021). The unicellular species had the lowest indel mutation rate, from 0.02 × 10^-9^ in yeast to 0.26 × 10^-9^ in algae, followed by arthropods with 0.55 (0.08, 1.24) × 10 . Mammals had a relatively higher indel mutation rate with 1.13 (0.43, 2.05) × 10, with humans at the higher end of this range at 1.74 (1.16, 2.50) × 10 . Of all the studied species, plants had the highest indel mutation rate, which was as high as 2.38 (0.51, 4.96) × 10.

**Figure 4.**
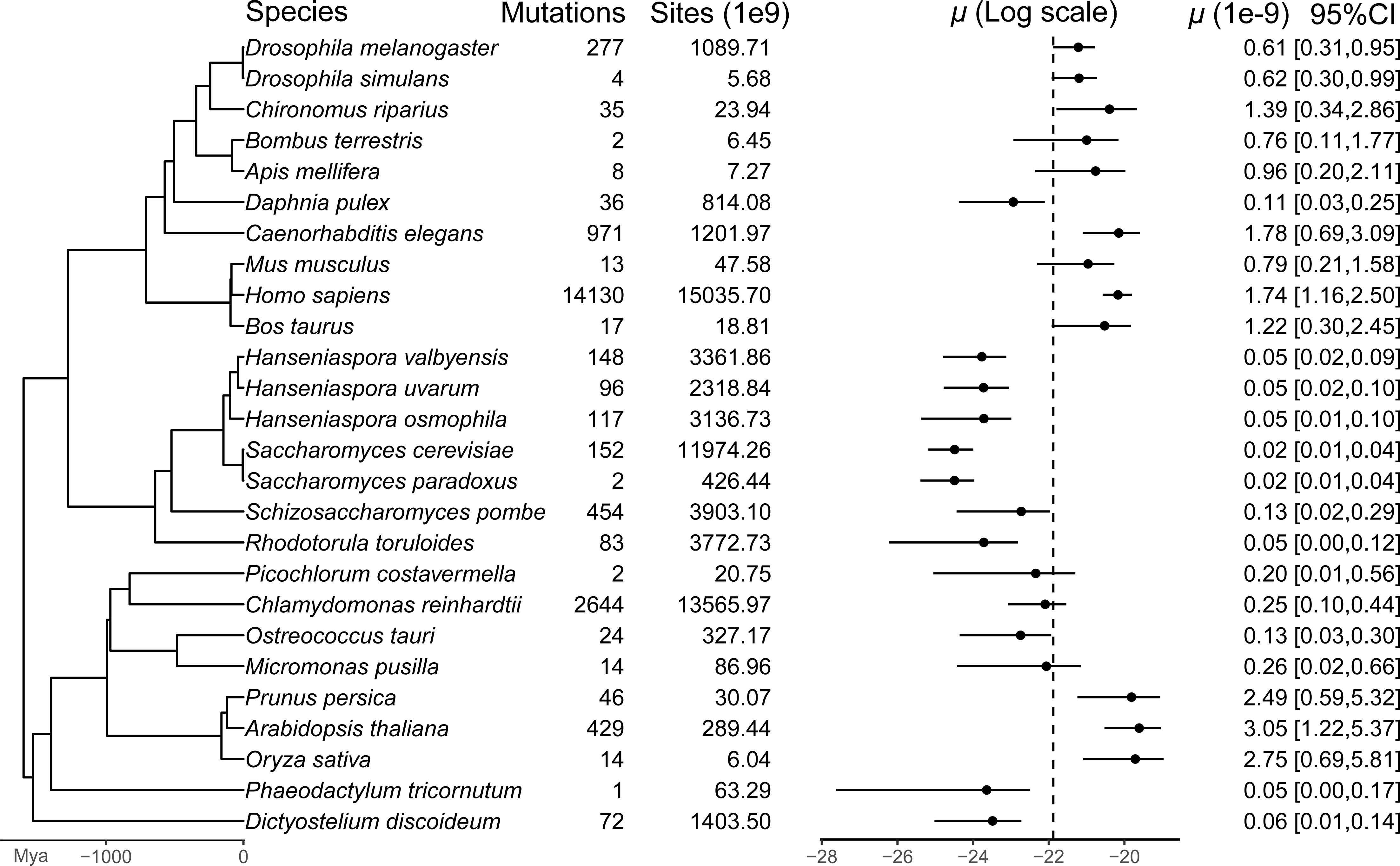
An analysis of short indel mutation rates for different species. The estimated mutation rates are presented on a log scale for clarity.

**Figure 5.**
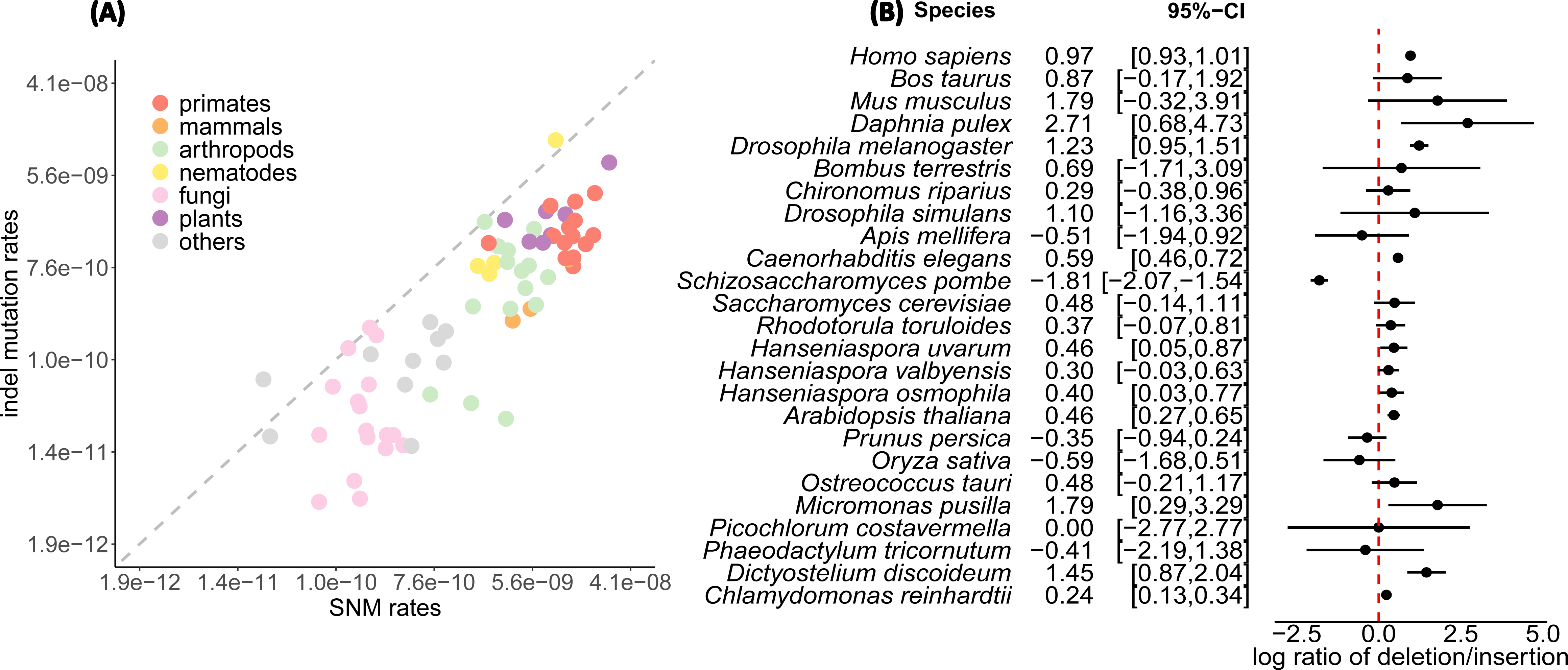
The correlation between indel mutation rates and SNM rates plotted on a log scale with a 1:1 ratio shown as a dashed line (A). The aggregated log ratios of deletion/insertion in eukaryotes (B).

Short deletions generally outnumbered short insertions across eukaryotes (Fig. 5B, Fig.S3). The ratio of deletions over insertions was as high as 15.0 in the water flea, 6.0 in mouse, 3.4 in D. melanogaster, 2.6 in humans, 1.8 in C. elegans, and 1.6 in Arabidopsis. Honeybees, diatoms and two plants (rice and peach) exhibited more insertions than deletions, but the difference wasn’t statistically significant (P > 0.05, binomial test). Fission yeast showed significantly more insertions overall.

A bias toward deletion events has been known for at least 40 years, since de Jong et al. (1981) found a four-fold excess of deletions over insertions in protein sequences. Petrov (2002) argued that the deletion bias was likely due to a thermodynamic disparity between short deletions and insertions that makes deletion events easier. However, studies in plants have suggested that unequal homologous recombination and illegitimate recombination may be another important cause of small deletions (Bennetzen et al., 2005). Whatever the cause, the dominance of deletions over insertions seems convincing, and many early studies argued that these short deletions resulted in genome contraction (Gregory, 2003; Petrov, 2002; Thomas et al., 2003). However, the role of deletions in shaping genome size remains controversial (Gregory, 2003, 2004). A recent study in birds and mammals indicated that short deletions alone cannot explain the DNA content loss (Kapusta et al., 2017), and we fail to find any evidence of a relationship between deletion rate and genome size here (Fig. S4; but note that power is likely to be low). It seems probable that other structural mutations, such as large deletions, gene duplication and transposable elements, may have greater impacts on genome size. For example, in Drosophila Wang et al. (2023) found transposable element insertions to be more frequent than SNMs, and an order of magnitude more frequent than short indels. Together, these studies suggest that the impact of short deletions on genome size is likely to be small, despite their higher frequency relative to short insertions.

### Concluding remarks

All genetic variation arises from de novo mutations. To estimate the rate at which these arise, studies have moved from investigating a few genes that result in phenotypic change, to whole genome sequencing of MA lines or families. Ten years ago, Campbell et al. (2013) called for more efforts to sequence genomes from non-human primate families to understand ‘how the rate has changed in different lineages’. And, in the present data-rich and tool-rich era we have now seen the sequencing of not just multiple non-human primates, but many taxa representing major clades across the tree of life. Nevertheless, the ‘tree of mutations’ is far from complete, at either the large or small scale. Even within animals, the vast majority of phyla have never been examined, and with the exception of primates, no large groups of very close relatives have been examined. Even for vertebrates—and despite of the recent large study by Bergeron et al. (2023) who examined 68 species–data are still scarce for reptiles and entirely lacking for amphibians.

However, as progress is being made toward a tree of mutation rates, attention still needs to focus on methodological heterogeneity. Different data filtering strategies may result in non-negligible differences in mutation rate estimates (Bergeron et al., 2022). Filters that are too conservative may discard some true mutations, while relaxed filters may inflate the false positive rate. Although many strategies have been proposed (Bergeron et al., 2022), no standardised pipeline has yet been agreed upon or widely adopted. All studies must therefore carefully address the issues of false positives by manual curation and/or PCR follow-up, and false negatives by simulation. Here, we did not examine the impact of the false positive rate or false negative rate, but we remind the readers of the potential impact from methodological difference, and caution that attention should be paid to the methods or parameters used in each study when comparisons are made.

Another potential source of bias may also come from the specific choice of samples used in each study. Beside the biological factors that affect mutation rates (e.g. age, sex), environmental factors can also influence mutation rates, for example temperature (Belfield et al., 2021; Waldvogel et al., 2021), osmotic stress (Hasan et al., 2022), or ultraviolet light (Xu et al., 2019). Samples collected from the wild rather than reared in laboratories may better reflect the ‘real’ mutation rate, but the variation that this introduces may also make it harder to draw broad conclusions. Despite these challenges, the data to come are exciting. In particular, as sequencing technology advances, we expect improved accuracy in long-read sequencing facilitating the phasing of mutations and permitting the easy detection of complex mutations and larger-scale structural changes.

## Methods

We made an exhaustive literature search for publications on the experimental estimation of mutation rate. We first searched Clarivate Web of Science up to the 21 September 2022 for ‘Title/Keywords/Abstract’ fields containing (“mutation rate” | “mutation rates” | “mutational rate” | “mutational rates”). As much of the resulting literature was related to somatic mutations (e.g., cancers) or virus mutations, we excluded ‘(“tumor” | “cancer” | “clinical” | “virus”)’ in the search field of ‘Title/Keywords/Abstract’. This search strategy resulted in 9462 studies with the earliest one dated 1928. We also searched for references to mutation rate estimates in other papers not captured by our search. As bacteria are outside of our study, we manually removed studies related to bacterial mutations. These filters resulted in 174 studies. Of these studies, we further excluded 20 studies that used phylogenetic approaches to infer mutation rates, 14 studies that used phenotypic markers, and two other studies that used recombination landscape or the site frequency spectrum to infer mutation rates statistically. In addition, we included one more recent study published after our initial submission (Bergeron et al., 2023), which estimated the mutation rates for 68 vertebrates using PO sequencing. This resulted in a total of 140 studies in our final dataset.

From each of the potential studies we extracted the year of the publication, the species, the population, reproductive mechanism (sexual, asexual), mutation type (SNM, short insertion, short deletion), the units of mutation rate (per bp per year, per bp per generation), the mean and 95% CI of mutation rate, the number of identified mutation events, the number of callable sites, the methods used to infer mutation rate (PO, MA or others), the sample size, the number of generations (if MA), the sequencing technology, the sequencing depth and other pertinent observations. Note that for multi-generation studies, such as MA, the number of ‘callable sites’ incorporates the possibility that a mutation may have arisen in any of several generations. If a 95% confidence or credible interval was not provided in the original study, we calculated a 95% confidence interval using ‘binconf()’ from the Hmisc R package (version 4.7-0), based on the number of mutation events and callable sites and assuming a binomial distribution (Girard et al., 2011). If a study did not provide the mutation count, we inferred an effective number of mutation events based on the reported mutation rate and its confidence interval. For studies in which the number of callable sites was not reported, an estimate was made from the number of mutation events and the reported mutation rate. If a study only reported the mutation rate, we assumed one mutation event and calculated the corresponding callable sites based on mutation rates. This latter approach will necessarily underestimate the precision with which rates are estimated (minimising their weight in the analysis), while keeping the estimate in the meta-analysis with unchanged mean values.

To compare the SNM rates across different species, we conducted a phylogenetic meta-analysis (i.e., a combined analysis of published data) using the mutation data from MA or PO studies. We obtained the phylogenetic relationships and relative divergence dates from http://www.timetree.org/ (Kumar et al., 2022) and visualised using ggtree (Yu, 2020). We performed the analyses using a phylogenetic generalized linear mixed model with a unit-scaled phylogeny as random effect, assuming a Poisson distribution for the occurrence of de novo mutations, off-set by the number of callable sites (i.e. enforcing a directly proportional relationship between the expected number of mutations and the number of sites observed). The analysis was conducted using the Bayesian mixed-model R-package MCMCglmm (version 2.32) (Hadfield, 2010) (see Supporting Materials). To investigate the effects of methods (MA or PO) used to estimate mutation rate, we additionally included ‘Method’ as an additional fixed effect in the above model.

We also investigated the relationships between mutation rate and generation time and genome size using univariate phylogenetic mixed models that included these factors separately as fixed effects, and a phylogenetic species term as a random effect. A log-transformation was made on these predictors before fitting the model (see Supporting Materials), and model predictions were made conditioning on each of the different species as random effect levels. The estimated generation time for each species was gathered from the literature (see Availability of data and materials) and we obtained estimates for 116 species. Genome sizes were taken to be the genome assembly size and gathered from the NCBI database, which resulted in 108 species with genome size available.

To test for a relationship between nucleotide diversity (π) and the per generation per-site SNM rate (µ) while accounting for uncertainty in both, we used MCMCglmm to fit a bivariate PGLMM with mutation number and π as responses, again including the number of callable sites as an offset on the number of mutations and generation time as a fixed effect (see Supporting Materials). We collected nucleotide diversity data for 102 species from Buffalo (2021), Bergeron et al. (2023) or literature (see Availability of data and materials).

## AVAILABILITY OF DATA AND MATERIALS

The datasets and the code to analyse the data are available on GitHub: https://github.com/Yiguan/mutation_literature

## ACKNOWLEDGEMENTS

We thank the authors of the many studies we analyse for making their data available, and particularly Lucie Bergeron and co-authors who generously shared recently published nucleotide diversity data with us. We thank Jarrod Hadfield for suggesting the overall statistical approach, helping with MCMCglmm syntax and interpretation, and drafting much of the supporting material. This work was funded by the UK Biotechnology and Biological Sciences Research Council through grant BB/T007516/1. For the purpose of open access, the authors have applied a Creative Commons Attribution (CC BY) licence to any Author Accepted Manuscript version arising from this submission.

## AUTHOR CONTRIBUTIONS

YW participated in data collection, performed analyses, and wrote the first draft. DJO wrote a part of the first draft and made extensive contributions to the development, drafting, and editing of the manuscript and supporting material, as well as model explanation. All authors gave final approval for publication.

## Figure legends

## Abbreviations

MA: mutation accumulation experiment
PO: parent-offspring experiment
SNMs: single nucleotide mutations
indels: insertions and deletion mutations
PGLMM: phylogenetic generalised linear mixed model
Mb: Mega base pair(s)
HPD: highest posterior density

## References

1. Baer, C. F., Miyamoto, M. M., & Denver, D. R. (2007). Mutation rate variation in multicellular eukaryotes: causes and consequences. Nature Reviews Genetics, 8(8), 619–631. doi:10.1038/nrg2158

2. Bailey, W. J., Fitch, D., Tagle, D. A., Czelusniak, J., Slightom, J. L., & Goodman, M. (1991). Molecular evolution of the psi eta-globin gene locus: gibbon phylogeny and the hominoid slowdown. Molecular Biology and Evolution, 8(2), 155–184.

3. Belfield, E. J., Brown, C., Ding, Z. J., Chapman, L., Luo, M., Hinde, E., Zheng, S. J. (2021). Thermal stress accelerates Arabidopsis thaliana mutation rate. Genome Research, 31(1), 40–50.

4. Bennetzen, J. L., Ma, J., & Devos, K. M. (2005). Mechanisms of recent genome size variation in flowering plants. Annals of Botany, 95(1), 127–132.

5. Bergeron, L. A., Besenbacher, S., Turner, T., Versoza, C. J., Wang, R. J., Price, A. L., Schierup, M. H. (2022). The Mutationathon highlights the importance of reaching standardization in estimates of pedigree-based germline mutation rates. Elife, 11. doi:10.7554/eLife.73577

6. Bergeron, L. A., Besenbacher, S., Zheng, J., Li, P., Bertelsen, M. F., Quintard, B., Shao, C. (2023). Evolution of the germline mutation rate across vertebrates. Nature, 1-7.

7. Bourguignon, T., Kinjo, Y., Villa-Martin, P., Coleman, N. V., Tang, Q., Arab, D. A., Ohkuma, M. (2020). Increased mutation rate is linked to genome reduction in prokaryotes. Current Biology, 30(19), 3848–3855. e3844.

8. Buffalo, V. (2021). Quantifying the relationship between genetic diversity and population size suggests natural selection cannot explain Lewontin’s Paradox. Elife, 10, e67509.

9. Burian, A. (2021). Does shoot apical meristem function as the germline in safeguarding against excess of mutations? Frontiers in Plant Science, 12, 707740.

10. Campbell, C. D., & Eichler, E. E. (2013). Properties and rates of germline mutations in humans. Trends in Genetics, 29(10), 575–584.

11. Chan, A. H., Jenkins, P. A., & Song, Y. S. (2012). Genome-wide fine-scale recombination rate variation in Drosophila melanogaster. Plos Genetics, 8(12), e1003090. doi:10.1371/journal.pgen.1003090

12. Chintalapati, M., & Moorjani, P. (2020). Evolution of the mutation rate across primates. Current Opinion in Genetics & Development, 62, 58–64.

13. Cruzan, M. B., Streisfeld, M. A., & Schwoch, J. A. (2022). Fitness effects of somatic mutations accumulating during vegetative growth. Evolutionary Ecology, 36(5), 767–785.

14. de Jong, W. W., & Rydén, L. (1981). Causes of more frequent deletions than insertions in mutations and protein evolution. Nature, 290(5802), 157–159.

15. Denver, D. R., Morris, K., Lynch, M., & Thomas, W. K. (2004). High mutation rate and predominance of insertions in the Caenorhabditis elegans nuclear genome. Nature, 430(7000), 679–682. doi:10.1038/nature02697

16. Drake, J. W. (1991). A constant rate of spontaneous mutation in DNA-based microbes. Proceedings of the National Academy of Sciences, 88(16), 7160–7164.

17. Drake, J. W., Charlesworth, B., Charlesworth, D., & Crow, J. F. (1998). Rates of spontaneous mutation. Genetics, 148(4), 1667–1686.

18. Drost, J. B., & Lee, W. R. (1995). Biological basis of germline mutation: comparisons of spontaneous germline mutation rates among drosophila, mouse, and human. Environmental and Molecular Mutagenesis, 25(S2), 48–64.

19. Freckleton, R. (2009). The seven deadly sins of comparative analysis. Journal of Evolutionary Biology, 22(7), 1367–1375.

20. Gao, Z., Wyman, M. J., Sella, G., & Przeworski, M. (2016). Interpreting the dependence of mutation rates on age and time. Plos Biology, 14(1), e1002355.

21. Girard, S. L., Gauthier, J., Noreau, A., Xiong, L., Zhou, S., Jouan, L., Rouleau, G. A. (2011). Increased exonic de novo mutation rate in individuals with schizophrenia. Nat Genet, 43(9), 860–863. doi:10.1038/ng.886

22. Gregory, T. R. (2003). Is small indel bias a determinant of genome size? Trends in Genetics, 19(9), 485–488.

23. Gregory, T. R. (2004). Insertion–deletion biases and the evolution of genome size. Gene, 324, 15–34.

24. Haag-Liautard, C., Dorris, M., Maside, X., Macaskill, S., Halligan, D. L., Houle, D., Keightley, P. D. (2007). Direct estimation of per nucleotide and genomic deleterious mutation rates in Drosophila. Nature, 445(7123), 82–85. doi:10.1038/nature05388

25. Hadfield, J. (2010). MCMC Methods for Multi-Response Generalized Linear Mixed Models: The MCMCglmm R Package. Journal of Statistical Software, 33(2), 1–22. doi:DOI 10.18637/jss.v033.i02

26. Hadfield, J. D., & Nakagawa, S. (2010). General quantitative genetic methods for comparative biology: phylogenies, taxonomies and multi-trait models for continuous and categorical characters. Journal of Evolutionary Biology, 23(3), 494–508. doi:10.1111/j.1420-9101.2009.01915.x

27. Halligan, D. L., & Keightley, P. D. (2009). Spontaneous mutation accumulation studies in evolutionary genetics. Annual Review of Ecology, Evolution, and Systematics, 40, 151–172.

28. Halliwell, B., Yates, L. A., & Holland, B. R. (2022). Multi-Response Phylogenetic Mixed Models: Concepts and Application. bioRxiv, 2022.2012. 2013.520338.

29. Harland, C., Charlier, C., Karim, L., Cambisano, N., Deckers, M., Mni, M., Georges, M. (2017). Frequency of mosaicism points towards mutation-prone early cleavage cell divisions in cattle. bioRxiv, 079863.

30. Harmon, L. J., Pennell, M. W., Henao-Diaz, L. F., Rolland, J., Sipley, B. N., & Uyeda, J. C. (2021). Causes and consequences of apparent timescaling across all estimated evolutionary rates. Annual Review of Ecology, Evolution, and Systematics, 52, 587–609.

31. Hasan, A. R., Lachapelle, J., El-Shawa, S. A., Potjewyd, R., Ford, S. A., & Ness, R. W. (2022). Salt stress alters the spectrum of de novo mutation available to selection during experimental adaptation of Chlamydomonas reinhardtii. Evolution.

32. Ho, E. K. H., Macrae, F., Latta, L. C., McIlroy, P., Ebert, D., Fields, P. D., Schaack, S. (2020). High and Highly Variable Spontaneous Mutation Rates in Daphnia. Molecular Biology and Evolution, 37(11), 3258–3266. doi:10.1093/molbev/msaa142

33. Ho, S. Y., Duchêne, S., & Duchêne, D. (2015). Simulating and detecting autocorrelation of molecular evolutionary rates among lineages. Molecular Ecology Resources, 15(4), 688–696.

34. Huang, W., Lyman, R. F., Lyman, R. A., Carbone, M. A., Harbison, S. T., Magwire, M. M., & Mackay, T. F. C. (2016). Spontaneous mutations and the origin and maintenance of quantitative genetic variation. Elife, 5. doi:10.7554/eLife.14625

35. Kaplanis, J., Ide, B., Sanghvi, R., Neville, M., Danecek, P., Coorens, T., McRae, J. (2022). Genetic and chemotherapeutic influences on germline hypermutation. Nature, 605(7910), 503–508.

36. Kapusta, A., Suh, A., & Feschotte, C. (2017). Dynamics of genome size evolution in birds and mammals. Proceedings of the National Academy of Sciences, 114(8), E1460–E1469.

37. Katju, V., & Bergthorsson, U. (2019). Old trade, new tricks: insights into the spontaneous mutation process from the partnering of classical mutation accumulation experiments with high-throughput genomic approaches. Genome biology and evolution, 11(1), 136–165.

38. Keightley, P. D., & Eyre-Walker, A. (2000). Deleterious mutations and the evolution of sex. Science, 290(5490), 331–333.

39. Keightley, P. D., Ness, R. W., Halligan, D. L., & Haddrill, P. R. (2014). Estimation of the Spontaneous Mutation Rate per Nucleotide Site in a Drosophila melanogaster Full-Sib Family. Genetics, 196(1), 313-+. doi:10.1534/genetics.113.158758

40. Keightley, P. D., Trivedi, U., Thomson, M., Oliver, F., Kumar, S., & Blaxter, M. L. (2009). Analysis of the genome sequences of three Drosophila melanogaster spontaneous mutation accumulation lines. Genome Research, 19(7), 1195–1201. doi:10.1101/gr.091231.109

41. Keith, N., Jackson, C. E., Glaholt, S. P., Young, K., Lynch, M., & Shaw, J. R. (2021). Genome-Wide Analysis of Cadmium-Induced, Germline Mutations in a Long-Term Daphnia pulex Mutation-Accumulation Experiment. Environmental Health Perspectives, 129(10). doi:10.1289/Ehp8932

42. Kimura, M. (1967). On the evolutionary adjustment of spontaneous mutation rates. Genetics Research, 9(1), 23–34.

43. Kimura, M. (1968). Evolutionary rate at the molecular level. Nature, 217(5129), 624–626.

44. Kondrashov, A. S. (2003). Direct estimates of human per nucleotide mutation rates at 20 loci causing Mendelian diseases. Human Mutation, 21(1), 12–27. doi:10.1002/humu.10147

45. Kondrashov, A. S., & Crow, J. F. (1993). A molecular approach to estimating the human deleterious mutation rate. Human Mutation, 2(3), 229–234. doi:10.1002/humu.1380020312

46. Kong, A., Frigge, M. L., Masson, G., Besenbacher, S., Sulem, P., Magnusson, G., Stefansson, K. (2012). Rate of de novo mutations and the importance of father’s age to disease risk. Nature, 488(7412), 471–475. doi:10.1038/nature11396

47. Konrad, A., Brady, M. J., Bergthorsson, U., & Katju, V. (2019). Mutational Landscape of Spontaneous Base Substitutions and Small Indels in Experimental Caenorhabditis elegans Populations of Differing Size. Genetics, 212(3), 837–854. doi:10.1534/genetics.119.302054

48. Kumar, S., Suleski, M., Craig, J. M., Kasprowicz, A. E., Sanderford, M., Li, M., Hedges, S. B. (2022). TimeTree 5: An expanded resource for species divergence times. Molecular Biology and Evolution, 39(8), msac174.

49. Lanfear, R. (2018). Do plants have a segregated germline? Plos Biology, 16(5), e2005439.

50. Li, W.-H., Tanimura, M., & Sharp, P. M. (1987). An evaluation of the molecular clock hypothesis using mammalian DNA sequences. Journal of Molecular Evolution, 25(4), 330–342.

51. Liu, H., & Zhang, J. (2021). The rate and molecular spectrum of mutation are selectively maintained in yeast. Nature Communications, 12(1), 1–11.

52. Lopez-Cortegano, E., Craig, R. J., Chebib, J., Samuels, T., Morgan, A. D., Kraemer, S. A., Keightley, P. D. (2021). De Novo Mutation Rate Variation and Its Determinants in Chlamydomonas. Molecular Biology and Evolution, 38(9), 3709–3723. doi:10.1093/molbev/msab140

53. Lynch, M. (1991). Methods for the analysis of comparative data in evolutionary biology. Evolution, 45(5), 1065–1080.

54. Lynch, M. (2008). The cellular, developmental and population-genetic determinants of mutation-rate evolution. Genetics, 180(2), 933–943.

55. Lynch, M. (2010). Evolution of the mutation rate. Trends in Genetics, 26(8), 345–352.

56. Lynch, M., & Walsh, B. (1998). Genetics and analysis of quantitative traits (Vol. 1): Sinauer Sunderland, MA.

57. Marais, G. A., Batut, B., & Daubin, V. (2020). Genome evolution: mutation is the main driver of genome size in prokaryotes. Current Biology, 30(19), R1083–R1085.

58. Martin, A. P., & Palumbi, S. R. (1993). Body size, metabolic rate, generation time, and the molecular clock. Proceedings of the National Academy of Sciences, 90(9), 4087–4091.

59. Meier, B., Cooke, S. L., Weiss, J., Bailly, A. P., Alexandrov, L. B., Marshall, J., Campbell, P. J. (2014). C. elegans whole-genome sequencing reveals mutational signatures related to carcinogens and DNA repair deficiency. Genome Research, 24(10), 1624–1636. doi:10.1101/gr.175547.114

60. Mukai, T., & Cockerham, C. C. (1977). Spontaneous mutation rates at enzyme loci in Drosophila melanogaster. Proceedings of the National Academy of Sciences, 74(6), 2514–2517.

61. Nachman, M. W., & Crowell, S. L. (2000). Estimate of the mutation rate per nucleotide in humans. Genetics, 156(1), 297–304.

62. Ness, R. W., Morgan, A. D., Colegrave, N., & Keightley, P. D. (2012). Estimate of the Spontaneous Mutation Rate in Chlamydomonas reinhardtii. Genetics, 192(4), 1447-+. doi:10.1534/genetics.112.145078

63. Nute, P. E., & Stamatoyannopoulos, G. (1984). Estimating Mutation-Rates Using Abnormal Human Hemoglobins. Yearbook of Physical Anthropology, 27, 135–151.

64. Obbard, D. J., Maclennan, J., Kim, K. W., Rambaut, A., O’Grady, P. M., & Jiggins, F. M. (2012). Estimating Divergence Dates and Substitution Rates in the Drosophila Phylogeny. Molecular Biology and Evolution, 29(11), 3459–3473. doi:10.1093/molbev/mss150

65. Ohno, M. (2019). Spontaneous de novo germline mutations in humans and mice: rates, spectra, causes and consequences. Genes & Genetic Systems, 94(1), 13–22.

66. Peck, K. M., & Lauring, A. S. (2018). Complexities of viral mutation rates. Journal of Virology, 92(14), e01031–01017.

67. Petrov, D. A. (2002). Mutational equilibrium model of genome size evolution. Theoretical Population Biology, 61(4), 531–544.

68. Rajaei, M., Saxena, A. S., Johnson, L. M., Snyder, M. C., Crombie, T. A., Tanny, R. E., Baer, C. F. (2021). Mutability of mononucleotide repeats, not oxidative stress, explains the discrepancy between laboratory-accumulated mutations and the natural allele-frequency spectrum in C. elegans. Genome Research, 31(9), 1602–1613.

69. Saxena, A. S., Salomon, M. P., Matsuba, C., Yeh, S. D., & Baer, C. F. (2019). Evolution of the Mutational Process under Relaxed Selection in Caenorhabditis elegans. Molecular Biology and Evolution, 36(2), 239–251. doi:10.1093/molbev/msy213

70. Schrider, D. R., Houle, D., Lynch, M., & Hahn, M. W. (2013). Rates and Genomic Consequences of Spontaneous Mutational Events in Drosophila melanogaster. Genetics, 194(4), 937-+. doi:10.1534/genetics.113.151670

71. Sigurðardóttir, S., Helgason, A., Gulcher, J. R., Stefansson, K., & Donnelly, P. (2000). The mutation rate in the human mtDNA control region. The American Journal of Human Genetics, 66(5), 1599–1609.

72. Stamatoyannopoulos, G., & Nute, P. E. (1982). De novo mutations producing unstable Hbs or Hbs M. II. Direct estimates of minimum nucleotide mutation rates in man. Human Genetics, 60(2), 181–188. doi:10.1007/BF00569709

73. Sung, W., Ackerman, M. S., Miller, S. F., Doak, T. G., & Lynch, M. (2012). Drift-barrier hypothesis and mutation-rate evolution. Proceedings of the National Academy of Sciences, 109(45), 18488–18492.

74. Thomas, G. W., & Hahn, M. W. (2014). The human mutation rate is increasing, even as it slows. Molecular Biology and Evolution, 31(2), 253–257.

75. Thomas, G. W., Wang, R. J., Puri, A., Harris, R. A., Raveendran, M., Hughes, D. S., Muzny, D. M. (2018). Reproductive longevity predicts mutation rates in primates. Current Biology, 28(19), 3193–3197. e3195.

76. Thomas, J. W., Touchman, J. W., Blakesley, R. W., Bouffard, G. G., Beckstrom-Sternberg, S. M., Margulies, E. H., Green, E. D. (2003). Comparative analyses of multi-species sequences from targeted genomic regions. Nature, 424(6950), 788–793. doi:10.1038/nature01858

77. Trimble, B. K., & Doughty, J. H. (1974). The amount of hereditary disease in human populations. Annals of Human Genetics, 38(2), 199–223. doi:10.1111/j.1469-1809.1974.tb01951.x

78. Waldvogel, A.-M., & Pfenninger, M. (2021). Temperature dependence of spontaneous mutation rates. Genome Research, 31(9), 1582-1589.

79. Wang, L., Ji, Y., Hu, Y., Hu, H., Jia, X., Jiang, M., Jia, Y. (2019). The architecture of intra-organism mutation rate variation in plants. Plos Biology, 17(4), e3000191.

80. Wang, R. J., Peña-Garcia, Y., Bibby, M. G., Raveendran, M., Harris, R. A., Jansen, H. T., Hahn, M. W. (2022). Examining the effects of hibernation on germline mutation rates in grizzly bears. Genome biology and evolution.

81. Wang, R. J., Thomas, G. W., Raveendran, M., Harris, R. A., Doddapaneni, H., Muzny, D. M., Hahn, M. W. (2020). Paternal age in rhesus macaques is positively associated with germline mutation accumulation but not with measures of offspring sociability. Genome Research, 30(6), 826–834.

82. Wang, Y., McNeil, P., Abdulazeez, R., Pascual, M., Johnston, S. E., Keightley, P. D., & Obbard, D. (2023). Variation in mutation, recombination, and transposition rates in Drosophila melanogaster and Drosophila simulans. Genome Research, gr. 277383.277122.

83. Wilding, C. (2017). Genetic diversity of the African malaria vector Anopheles gambiae. Nature, 552, 96–100.

84. Wu, C.-I., & Li, W.-H. (1985). Evidence for higher rates of nucleotide substitution in rodents than in man. Proceedings of the National Academy of Sciences, 82(6), 1741–1745.

85. Wu, F., & Przeworski, M. (2022). A paternal bias in germline mutation is widespread in amniotes and can arise independently of cell division numbers. Elife, 11, e80008–e80008.

86. Wu, F. L., Strand, A. I., Cox, L. A., Ober, C., Wall, J. D., Moorjani, P., & Przeworski, M. (2020). A comparison of humans and baboons suggests germline mutation rates do not track cell divisions. Plos Biology, 18(8), e3000838.

87. Xu, S., Stapley, J., Gablenz, S., Boyer, J., Appenroth, K. J., Sree, K. S., Huber, M. (2019). Low genetic variation is associated with low mutation rate in the giant duckweed. Nature Communications, 10(1), 1–6.

88. Yang, S. H., Wang, L., Huang, J., Zhang, X. H., Yuan, Y., Chen, J. Q., Tian, D. C. (2015). Parent-progeny sequencing indicates higher mutation rates in heterozygotes. Nature, 523(7561), 463–U187. doi:10.1038/nature14649

89. Yoder, A. D., & Tiley, G. P. (2021). The challenge and promise of estimating the de novo mutation rate from whole-genome comparisons among closely related individuals. Molecular Ecology, 30(23), 6087–6100. doi:10.1111/mec.16007

90. Yu, G. (2020). Using ggtree to visualize data on tree-like structures. Current protocols in bioinformatics, 69(1), e96.

