## Supporting Materials for "Experimental estimates of germline mutation rate in eukaryotes: a phylogenetic meta-analysis"

### Supporting Methods

#### 1 Introduction to PGLMMs and mutation rate

Generalised Linear Models (GLMs) assume that observations are drawn from a distribution whose mean, after a link-transformation, can be modelled as a linear combination of predictors with estimated weights ('parameters')(Hardin et al., 2007). Generalised Linear Mixed Models (GLMMs) extend this framework through the inclusion of *random effects*, that is, weights which are themselves random variables (Bolker et al., 2009).

In the context of multi-species analyses, species do not provide independent observations because patterns of shared ancestry cause observations of related species to be correlated (Freckleton, 2009). That is, *a priori*, more closely related species are expected to be less divergent in any trait than are distantly related species. This non-independence is accounted for in 'Phylogenetic' GLMMs, which model the covariance between species as being proprotional to the duration of their shared ancestry (Hadfield, 2010).

The occurrence of rare independent events, such as mutations, can usefully be modelled in a GLMM by assuming the observed number of events is drawn from a Poisson distribution, for which the natural link function is $log_e()$. That is, the GLMM predicts the $log_e()$ of the expected number of mutations. If the number of detected mutations scales linearly with the number of observations made (i.e. the number of callable sites) and $log_e$(Sites) is included as a predictor (or offset) then, since $log_e(E[Mutations]) = 1 \cdot log_e(Sites)$ implies $E[Mutations] = Sites^1$, the model prediction is the $log_e()$ of the expected number of mutations per site, i.e. the mutation rate.

#### 2 Univariate model for mutation rate

We first model the observed number of mutations in a univariate Phylogenetic Poisson Mixed Model in which the expected number of mutations is predicted by the genome size, generation time, number of callable sites, and species identity. Our full log-linear model for the number of mutations observed in experiment $j$ and species $i$ is thus:

$$\begin{aligned} log_e(E[Mutations_{ij}]) = \beta_0^{(M)} &+ log_e(Sites_{ij})\beta_1^{(M)} \\ &+ log_e(Gsize_i)\beta_2^{(M)} + log_e(Gtime_i)\beta_3^{(M)} \\ &+ u_i^{(M)} + e_{ij}^{(M)} \end{aligned} \tag{1}$$

where $\beta_0^{(M)}$ is the intercept, and the regression slope $\beta_1^{(M)}$ is fixed very close to one, so that $log_e(Sites_{ij})$ functions as an offset. Note that by taking logs of the predictors, we assume that a proportional change in the predictor induces a proportional change in the number of mutations. The term $u_i^{(M)}$ is a random species effect and $e_{ij}^{(M)}$ an observation-level random effect that accounts for any overdispersion (i.e. any remaining variation among Poisson expectations). The observation-level random effects are assumed to be independently and identically distributed normal variates with variance $\sigma^2_{e^{(M)}}$. The vector of species effects is assumed to follow a multivariate normal distribution with zero mean vector, and covariance $\sigma^2_{u^{(M)}}\mathbf{A}$, where $\mathbf{A}$ is a matrix of shared branch lengths and $\sigma^2_{u(M)}$ is the phylogenetic variance. The distribution of the number of mutations is then Poisson, with mean equal to the exponentiated log-linear model above.

We fitted this model in R, using the Bayesian mixed model fitting package 'MCMCglmm' (Hadfield, 2010), specifying broad weakly-informative priors on each of the parameters. We also fitted three smaller variations on this model, omitting either or both of the terms for generation time ($Gtime_i$) and genome size ($Gsize_i$). The syntax used to fit the full model was:

```
# Prior Specification
vv <- diag(c(1e6, 1e-8, 1e6, 1e6))
prior <- list(
                B=list(mu=c(0,1,0,0), V=vv),
                G=list(G1=list(V=1, nu=1, alpha.mu=0, alpha.V=100))
            )

# Model Fitting
MCMCglmm(   Mutations ~ log_callable + log_Gsize_Mb + log_Gtime_Year,
            random= ~ Species,
            data=ccz,
            ginverse=list(Species=InverseTree),
            prior = prior,
            family = 'poisson',
            nitt=1001000, thin=1000, burnin=10000,
            pr=TRUE, verbose = FALSE
        )
```

In the code above, the prior comprises a list of two elements: 'B', the prior for the fixed effects, and 'G', a prior for the random effect. The first list element, 'B', includes a vector of prior means for the model intercept (mean zero), callable sites (mean one), genome size (mean zero), and generation time (mean zero), along with a diagonal matrix of prior variances (covariances are set to zero). Note that to achieve the offset, the prior for the variance associated with the 'callable sites' slope parameter is set to a very small value, effectively fixing the gradient at one. The second list element, 'G', specifies a parameter-expanded prior for the phylogenetic variance that induces a scaled-$F_{1,1}$ prior on the variance with a scale of $\sqrt{100} = 10$ (Hadfield, 2014). Within the function call to MCMCglmm() the argument 'ginverse' supplies the inverse of the distance matrix that describes species relationships in an ultrametric tree that has been re-scaled to have a total tree depth of one. The arguments 'nitt', 'thin', and 'burnin' respectively control the length, sampling frequency, and initial un-sampled steps for the MCMC chain, and 'pr' specifies that estimates for random effects should be recorded.

#### 3 Multivariate model for diversity

The drift barrier hypothesis predicts that species with high genetic drift (i.e., small $N_e$) will tend to have a high per-site per-generation mutation rate $\mu$ (Lynch, 2010). Unfortunately, long-term $N_e$ must be inferred from pairwise 'neutral' (unconstrained) genetic diversity $\pi_s$ by making use of the relationship $N_e = \pi_s/4\mu$, and error in the estimates could drive a spurious relationship in a regression of $\mu$ on $\pi_s/4\mu$. Instead, we make use of a bivariate PGLMM that models the covariance between dependent variables $\mu$ and $\pi_s$, parameterised to estimate the gradient of the relationship (see below). We expect that $\pi_s = 4N_e\mu$ and so, on the $log_e$ scale of our analysis, $log_e(\pi_s) = log_e(4) + log_e(N_e) + log_e(\mu)$ such that the regression slope of $log_e(\pi)$ on $log_e(\mu)$ should be 1, unless $N_e$ (which is not measured) is correlated with $\mu$. If the gradient is less than 1, then this suggests that species with large $N_e$ have a lower mutation rate than species with small $N_e$. However, it should be noted that if $\pi_s$ is not neutral, for example there is weak constraint that has a greater impact in large populations, this too could depress the gradient below 1.

As only a single observation of $\pi_s$ is available per species, we chose to simplify the model for mutation rate by combining mutation counts (and callable sites) across experiments, so that there is also a single mutation rate for each species. We also chose to omit genome size as a predictor, as it was not significant in the full univariate model above. Our full log-linear bivariate model for the number of mutations and genetic diversity observed in species $i$ would therefore be:

$$\begin{aligned} log_e(E[Mutations_i]) &= \beta_0^{(M)} + log_e(Sites_i)\beta_1^{(M)} + log_e(Gtime_i)\beta_2^{(M)} + u_i^{(M)} + e_i^{(M)} \\ log_e(E[Diversity_i]) &= \beta_0^{(D)} + log_e(Gtime)\beta_1^{(D)} + u_i^{(D)} + e_i^{(D)} \end{aligned} \tag{2}$$

We can then model covariances between the phylogenetic effects of the two response variables as $COV(\mathbf{u}^{(D)}, \mathbf{u}^{(M)}) = \sigma_{u^{(D,M)}}\mathbf{A}$, where $\sigma_{u^{(D,M)}}$ is the phylogenetic covariance between log diversity and log mutation rate. We can also model covariances between the non-phylogenetic species-level random effects as $COV(\mathbf{e}^{(D)}, \mathbf{e}^{(M)}) = \sigma_{e^{(D,M)}}$. As before, $\beta_1^{(M)}$ is set to 1, and acts as an offset.

However, the equations in (2) above are parameterised in terms of the covariance between mutation rate and diversity, whereas we are are explicitly interested in the gradient of the regression slope. Usefully, a $k \times k$ unstructured covariance matrix can be re-parameterised in terms of a $(k-1)^{th}$ antedependence structure, and in the $k = 2$ case, one of the parameters is then the regression slope from a regression of one variable on the other ($u^{(D)}$ on $u^{(M)}$ in our case) (Thomson et al., 2017):

$$\mathbf{V}_u = \begin{bmatrix} \sigma^2_{u^{(M)}} & \sigma_{u^{(M,D)}} \\ \sigma_{u^{(D,M)}} & \sigma^2_{u^{(D)}} \end{bmatrix} = \begin{bmatrix} \sigma^2_{u^{(M)}} & (\beta^{(D|M)}\sigma_{u^{(M)}})^2 \\ (\beta^{(D|M)}\sigma_{u^{(M)}})^2 & (\beta^{(D|M)}\sigma_{u^{(M)}})^2 + \sigma^2_{u^{(D|M)}} \end{bmatrix} \tag{3}$$

where $\beta^{(D|M)}$ is the regression of $u^{(D)}$ on $u^{(M)}$ and $\sigma^2_{u^{(D|M)}}$ is the (residual) variance in $u^{(D)}$ after conditioning on $u^{(M)}$. We can then rewrite the full bivariate model as:

$$\begin{aligned} log_e(E[Mutations_i]) &= \beta_0^{(M)} + log_e(Sites_i)\beta_1^{(M)} + log_e(Gtime_i)\beta_2^{(M)} + u_i^{(M)} + e_i^{(M)} \\ log_e(E[Diversity_i]) &= \beta_1^{(D)} + log_e(Gtime)\beta_2^{(D)} + u_i^{(M)}\beta_u^{(D|M)} + u_i^{(D|M)} + e_i^{(M)}\beta_e^{(D|M)} + e_i^{(D|M)} \end{aligned} \tag{4}$$

where $\beta_u^{(D|M)}$ is the regression coefficient for the regression of diversity on mutation rate for the phylogenetic component of variation, and $\beta_e^{(D|M)}$ is the regression coefficient for the regression of diversity on mutation rate for the remaining (non-phylogenetic) components of variation. This re-parameterisation not only permits us direct access to the gradient of interest, but also allows a prior to be directly placed on that parameter. As before, we fitted this model in R, using the Bayesian mixed model fitting package 'MCMCglmm' (Hadfield, 2010). The syntax used to fit the full model was:

```
#Prior Specification
vv <- diag(5)
diag(vv) <- c(1e6, 1e6, 1e6, 1e6, 1e-8)
prior <- list(
                B=list(mu=c(0,0,0,0,1),V=vv),
                G=list(G1=list( V=diag(2), nu=2,
                        alpha.V=diag(2)*100, beta.mu=1, beta.V=10)),
                R=list( R=list(V=diag(2), nu=0.002, beta.mu=1, beta.V=10)
```

```
            )

#Model Fitting
MCMCglmm(   y ~ trait - 1
                + trait:log_Gt
                + at.level(trait, 'Mutations'):log_callable,
            random = ~ante1(trait):Species,
            rcov = ~ante1(trait):units,
            data = d12.y,
            ginverse = list(Species=InverseTree),
            prior = prior,
            nitt=1001000, thin=1000, burnin=10000,
            family = NULL, verbose = FALSE
        )
```

In the code above, a single data-frame column 'y' contains both the counts of mutations and the estimates of genetic diversity. These are indexed by the column 'trait', and the term '-1' allows these two 'traits' to have different intercepts. The term 'at.level()' is used to apply the offset only to the 'Mutations' trait. The argument 'random = ∼ ante1(trait):Species' specifies the antedependence structure for the phylogenetic covariance matrix, and the argument 'rcov = ∼ ante1(trait):units' specifies the antedependence structure for the remaining non-phylogenetic (residual) covariance matrix. Note that the distributions for the 'traits' Mutations and Diversity (Poisson and Gaussian, respectively) were specified by an additional column 'family' in the data-frame, rather than by using the argument 'family=', which is set to 'NULL'. As in the univariate case above, the gradient relating the mutation rate to the number of 'callable sites' is fixed at 1 using a prior mean of 1 with a very small variance. As before, the random (phylogenetic) effect has broad weakly-informative parameter expanded priors. However, as we wish to test whether the gradient is different from 1, we conservatively set the prior mean for the gradient at 1, rather than zero.

### References


B. M. Bolker, M. E. Brooks, C. J. Clark, S. W. Geange, J. R. Poulsen, M. H. H. Stevens, and J.-S. S. White. Generalized linear mixed models: a practical guide for ecology and evolution. *Trends in ecology & evolution*, 24(3):127–135, 2009.

R. Freckleton. The seven deadly sins of comparative analysis. *Journal of evolutionary biology*, 22(7):1367–1375, 2009.

J. Hadfield. Mcmc methods for multi-response generalized linear mixed models: the mcmcglmm r package. *Journal of statistical software*, 33:1–22, 2010.

J. Hadfield. Mcmcglmm course notes. *available at: cran. r-project. org/web/packages/MCMCglmm/vignettes/CourseNotes. pdf*, 2014.

J. W. Hardin, J. W. Hardin, J. M. Hilbe, and J. Hilbe. *Generalized linear models and extensions.* Stata press, 2007.

M. Lynch. Evolution of the mutation rate. *TRENDS in Genetics*, 26(8):345–352, 2010.

C. E. Thomson, F. Bayer, N. Crouch, S. Farrell, E. Heap, E. Mittell, M. Zurita-Cassinello, and J. D. Hadfield. Selection on parental performance opposes selection for larger body mass in a wild population of blue tits. *Evolution*, 71(3):716–732, 2017.
