## Supplementary figures and images for "Experimental estimates of germline mutation rate in eukaryotes: a phylogenetic meta-analysis"

### Fig. S1

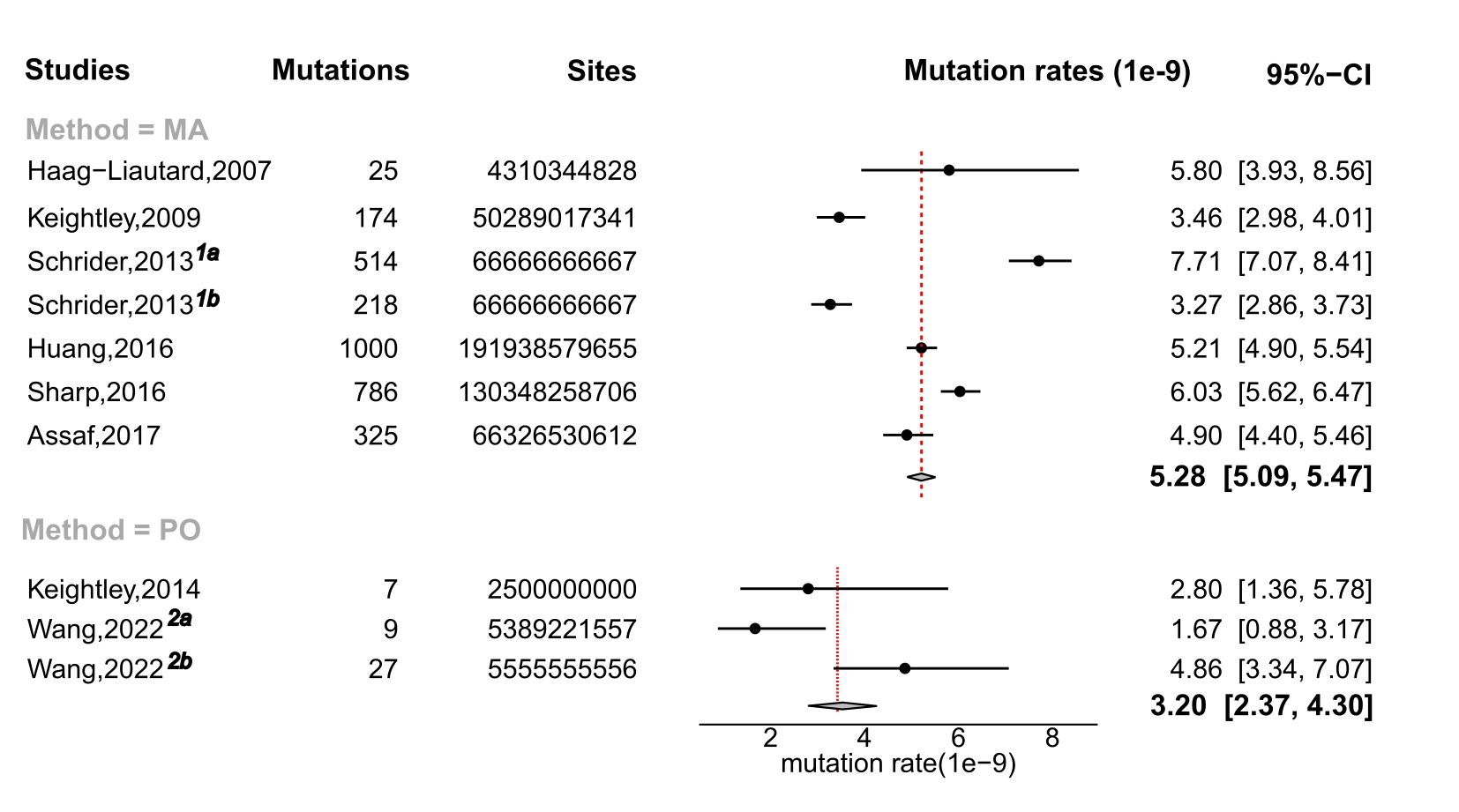

### Fig. S2

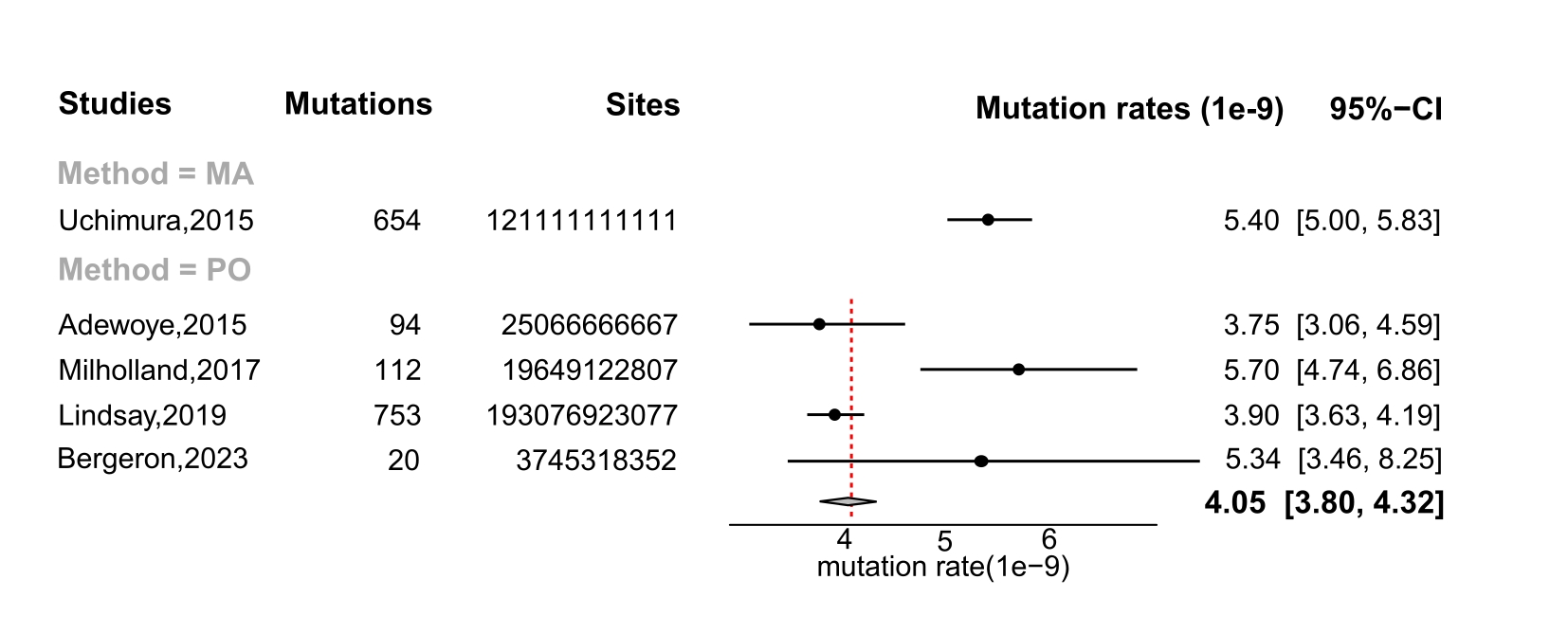

### Fig. S3

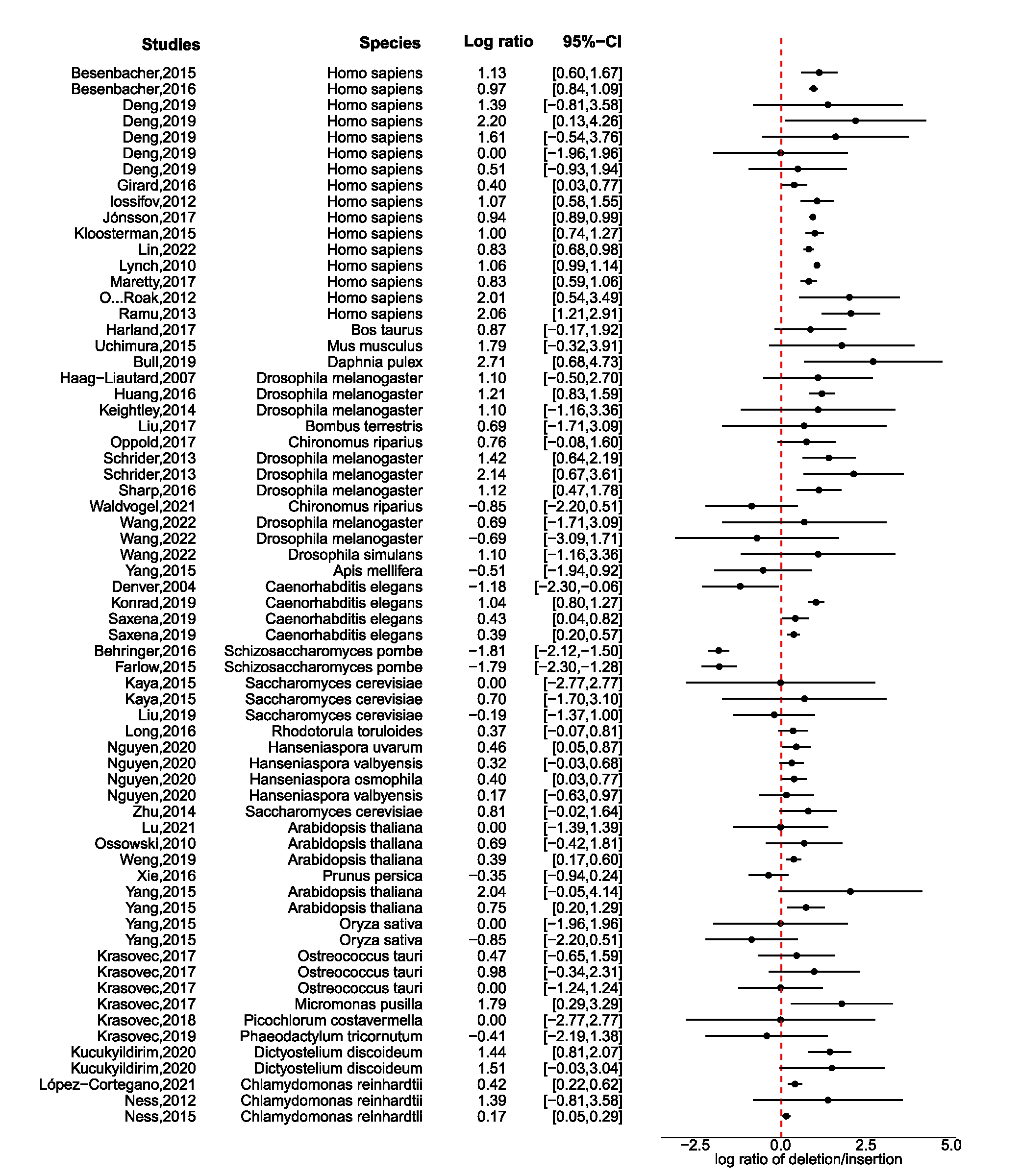

### Fig. S4

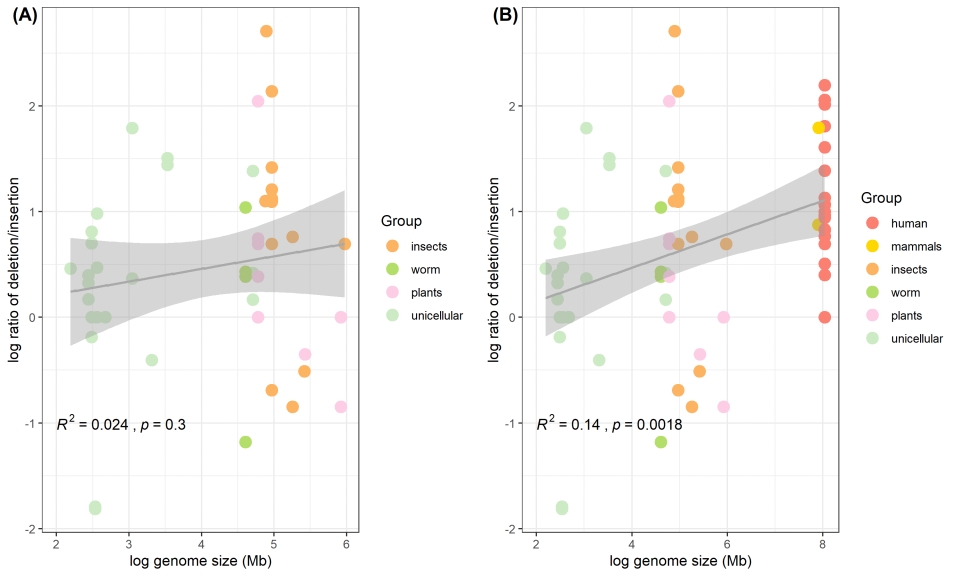
